# Hepatocyte TEAD1 drives epithelial-stromal remodeling during cholestatic liver injury

**DOI:** 10.64898/2026.05.21.726939

**Authors:** Amit Kumar, Jeongkyung Lee, Vinny Negi, Varun Mandi, Domenic Filingeri, Joey Danvers, Rajat Pant, Samit Ghosh, Mousumi Moulik, Vijay K Yechoor

**Affiliations:** Division of Endocrinology, Diabetes & Metabolism, Department of Medicine, University of Pittsburgh, PA, USA; Department of Medicine, Pittsburgh Heart, Lung, and Blood Vascular Medicine Institute, University of Pittsburgh, PA, USA; Division of Pediatric Cardiology, UPMC Children’s Hospital of Pittsburgh, University of Pittsburgh, PA, USA

## Abstract

**Background & Aims:** Primary sclerosing cholangitis (PSC) is a progressive cholangiopathy characterized by ductular remodeling, inflammation, and periportal fibrosis, for which effective medical therapies remain limited. The Hippo pathway effector TEAD1 has been implicated in liver regeneration and fibrogenesis; however, its role in cholestatic injury remains poorly defined. We investigated whether hepatocyte TEAD1 regulates injury-associated remodeling in a PSC-mimicking model and whether this mechanism is conserved in human PSC liver.

**Methods:** Hepatocyte-specific TEAD1 knockout mice (Alb-TEAD1^-/-^) and littermate controls were subjected to DDC-induced cholestatic injury. Ductular reaction, fibrosis, inflammation, and bile acid-related gene programs were assessed by histology, immunostaining, and gene expression analyses. Translational relevance was evaluated using bulk and single-cell transcriptomic datasets from human PSC liver.

**Results:** Hepatocyte TEAD1 deletion attenuated DDC-induced fibrosis, ductular expansion, and inflammatory cell accumulation, while preserving hepatocyte proliferative responses. TEAD1-deficient livers exhibited reduced expression of profibrotic mediators, including *Spp1, Ctgf*, and *Cyr61*, with decreased extracellular matrix deposition. In contrast, canonical transcriptional adaptations to cholestatic stress, including suppression of bile acid uptake, induction of efflux pathways, and repression of bile acid synthesis genes, were preserved in the absence of TEAD1.

Analysis of human PSC datasets demonstrated coordinated upregulation of TEAD1 and TEAD-associated target genes. Single-cell transcriptomic analysis further revealed hepatocyte-enriched TEAD1 expression and activation of a TEAD1 target gene program across all hepatic zones in PSC, with effect sizes exceeding those observed in non-parenchymal populations. TEAD1 activation was accompanied by co-expression of profibrotic mediators and downregulation of hepatocyte differentiation markers, consistent with a maladaptive hepatocyte state.

**Conclusions:** Hepatocyte TEAD1 drives ductular, inflammatory, and fibrogenic remodeling during cholestatic injury without disrupting bile acid metabolic adaptation. These findings identify TEAD1 as a hepatocyte-intrinsic regulator of epithelial-stromal crosstalk and establish conserved activation of this pathway in human PSC, supporting TEAD-directed signaling as a therapeutic target.

## INTRODUCTION

Liver fibrosis represents the cumulative outcome of chronic injury and sustained inflammatory signaling, characterized by progressive extracellular matrix deposition and architectural remodeling. This process reflects coordinated interactions among hepatocytes, cholangiocytes, immune cells, and hepatic stellate cells (HSCs), ultimately leading to activation of fibrogenic pathways and loss of tissue integrity. Across diverse etiologies, including metabolic dysfunction-associated steatohepatitis (MASH) and cholestatic liver diseases, persistent injury frequently progresses to advanced fibrosis and organ failure.

Primary sclerosing cholangitis (PSC) is a progressive cholangiopathy marked by chronic biliary injury, ductular reaction, and periportal fibrosis. Despite progress in understanding the disease pathogenesis, no effective disease-modifying therapies exist, and liver transplantation remains the only definitive treatment for advanced disease. The 3,5-diethoxycarbonyl-1,4-dihydrocollidine (DDC) diet-induced injury model recapitulates key features of cholestatic liver injury, including ductular expansion, biliary obstruction, and periductal fibrosis [1]. With sustained injury, impaired hepatocyte regenerative responses are accompanied by enhanced ductular signaling, a hallmark of chronic cholestatic remodeling [2]. These features highlight the role of hepatocyte-derived signals in coordinating epithelial-stromal interactions during cholestatic injury.

The transcriptional enhancer-associated domain protein 1 (TEAD1), a downstream effector of Hippo signaling, regulates cellular proliferation, mechano-transduction, and tissue remodeling. In the liver, TEAD1 activation has been associated with hepatocyte proliferation during regeneration and is increased in metabolic liver disease [3, 4]. In non-parenchymal compartments, TEAD1 regulates profibrotic transcriptional programs, including CTGF and CYR61, and promotes stellate cell activation and extracellular matrix production [5, 6]. However, the contribution of hepatocyte-intrinsic TEAD1 signaling to cholestatic injury and fibrogenic remodeling remains poorly defined.

In particular, whether hepatocyte-intrinsic TEAD1 signaling coordinates ductular remodeling, inflammatory activation, and fibrogenic responses during cholestatic injury remains unresolved. Here, we investigate the role of hepatic TEAD1 in a DDC-induced model of cholestatic injury. We demonstrate that hepatocyte TEAD1 deficiency attenuates ductular expansion, inflammation, and fibrogenic remodeling while preserving hepatocyte proliferative capacity, identifying TEAD1 as a regulator of injury-associated epithelial-stromal crosstalk. Extending these findings to human disease, single-cell transcriptomic analyses of PSC liver reveal hepatocyte-enriched TEAD1 expression and activation of a TEAD1-dependent transcriptional program across all hepatic zones. Collectively, these data position hepatocyte TEAD1 as a conserved driver of maladaptive remodeling in cholestatic liver disease and support TEAD1-directed pathways as potential therapeutic targets in PSC.

## RESULTS

### Hepatocyte TEAD1 is induced during cholestatic injury and efficiently deleted in Alb-Tead1 knockout livers

To investigate the role of hepatocyte TEAD1 in cholestatic liver injury, we utilized a DDC-induced injury model in hepatocyte-specific TEAD1 knockout (Alb-TEAD1^-/-^) mice (Figure 1A). Immunoblot analysis demonstrated induction of TEAD1 protein in control (TEAD1^fl/fl^) livers following DDC administration, whereas TEAD1 expression was markedly reduced in Alb-TEAD1^-/-^ livers (Figure 1B). Immunofluorescence analysis further showed reduced TEAD1 signal in albumin-positive hepatocytes in Alb-TEAD1^-/-^ livers compared with controls (Figure 1C), supporting effective hepatocyte-targeted deletion. Consistent with prior reports of systemic effects of DDC feeding, mice exhibited weight loss following DDC treatment; however, hepatocyte-specific deletion of TEAD1 did not significantly alter overall weight loss (Figure 1D). Liver weights were modestly increased in Alb-TEAD1^-/-^ mice following DDC exposure (Figure 1E). Together, these data demonstrate induction of TEAD1 during cholestatic injury and effective hepatocyte-specific deletion in Alb-TEAD1^-/-^ livers.

**Figure 1.**
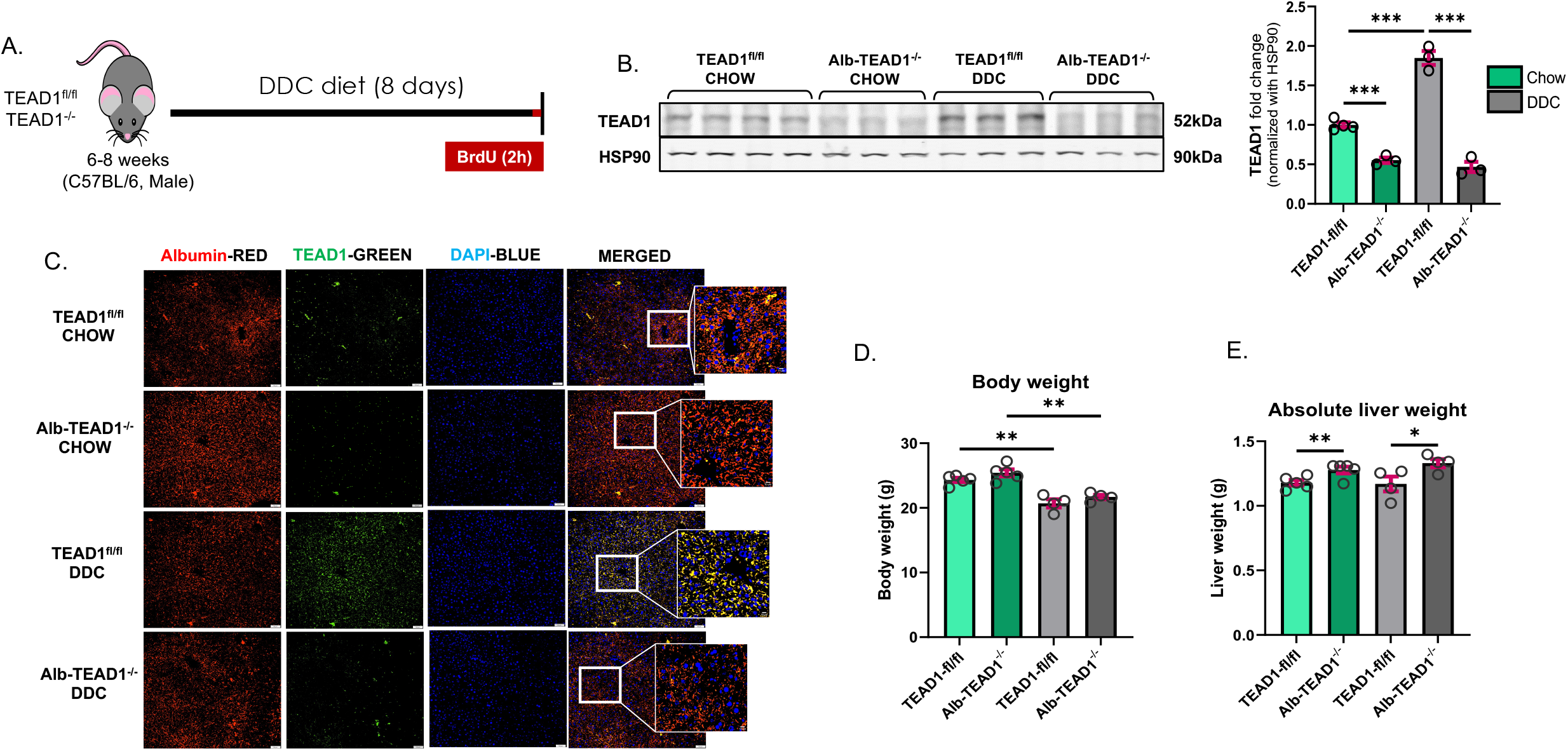
Induction and hepatocyte-specific deletion of TEAD1 in DDC-induced cholestatic liver injury. **(A)** Schematic of the experimental design-hepatocyte-specific TEAD1 knockout (Alb-TEAD1^-/-^) mice were generated by crossing TEAD1^fl/fl^ mice with Alb-Cre transgenic mice and subjected to 3,5-diethoxycarbonyl-1,4-dihydrocollidine (DDC) diet–induced cholestatic injury. **(B)**Representative immunoblot of TEAD1 protein from liver 30μg lysates of chow and DDC-fed TEAD1^fl/fl^ and Alb-TEAD1^-/-^ mice. HSP90 was used as a loading control. **(C)** Representative immunofluorescence images of Albumin (red) with TEAD1 (green) and nuclear counterstain (DAPI, blue) showing TEAD1 upregulation in TEAD1^fl^/^fl^ livers following DDC treatment and TEAD1 deletion in Alb-TEAD1^-/-^ hepatocytes. **(D)** Body weight following DDC treatment in control and Alb-TEAD1^-/-^ mice. **(E)** Absolute liver weight in control and Alb-TEAD1^-/-^ mice under chow and DDC conditions. Data are presented as mean ± SEM (n = 4-5 mice per group). Images were taken at 20X and insets show magnified view, scale bar = 10 µm. [Statistical significance was determined by one-way ANOVA with post hoc testing (*p ≤ 0.05, **p ≤ 0.01, ***p ≤ 0.001)].

### Hepatocyte TEAD1 deficiency attenuates ductular remodeling during DDC-induced injury

A prominent feature of cholestatic liver disease is the expansion of ductular structures in response to injury. Histological examination of DDC-treated livers revealed marked periportal remodeling and intraductal pigment accumulation in control mice, whereas these features were notably reduced in Alb-TEAD1^-/-^ livers (Figure 2A). To directly assess ductular expansion, we performed immunofluorescence staining for CK19, a marker of biliary epithelial cells and ductular reaction. DDC treatment induced robust CK19^+^ ductular expansion in control livers, whereas Alb-TEAD1^-/-^ livers exhibited a clear reduction in CK19^+^ structures (Figure 2B). These findings indicate that hepatocyte TEAD1 contributes to ductular remodeling in the setting of cholestatic injury.

**Figure 2.**
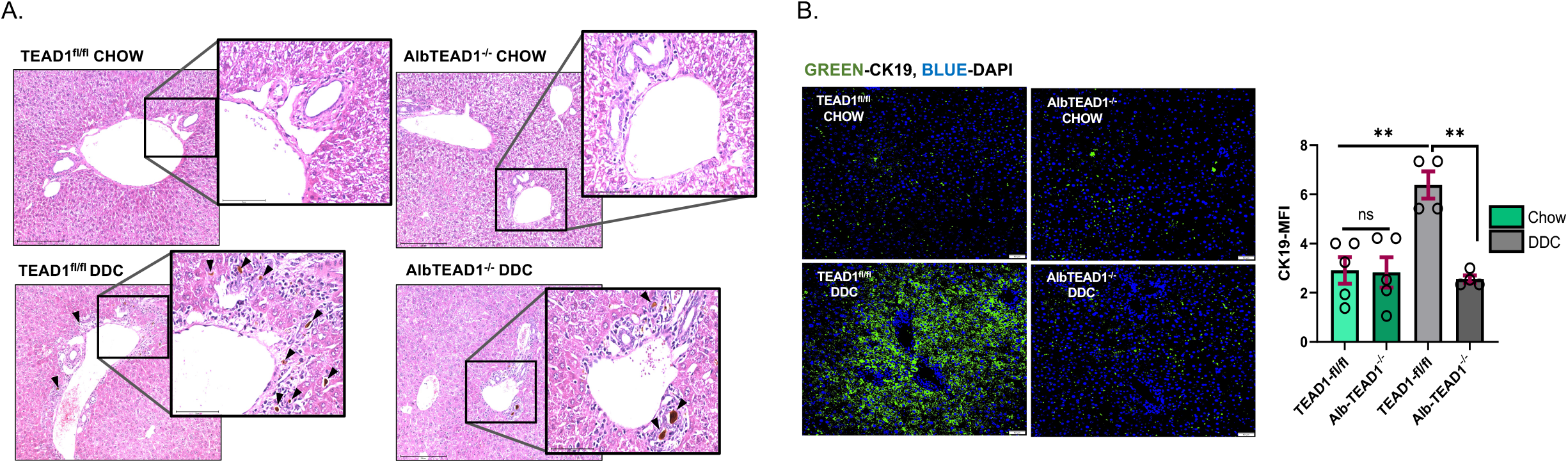
Reduced ductular expansion in Alb-TEAD1^-/-^ livers following DDC-induced injury. **(A)** Representative hematoxylin and eosin (H&E)–stained liver sections from chow and DDC-fed TEAD1^fl/fl^ and Alb-TEAD1^-/-^ mice. **(B)** Representative immunofluorescence images of CK19 (green) with nuclear counterstain (DAPI, blue) showing ductular reaction in control and Alb-TEAD1^-/-^ livers following DDC treatment. Data are presented as mean ± SEM (n = 3-5 mice per group). Images were taken at 20X and insets show a 60X magnified view. [Statistical significance was determined by one-way ANOVA with post hoc testing (**p ≤ 0.01)].

### Hepatocyte TEAD1 promotes fibrogenic remodeling and profibrotic gene expression

Given the close association between ductular reaction and fibrogenesis, we next examined whether hepatocyte TEAD1 influences fibrotic remodeling. Picro-Sirius Red staining demonstrated substantial periportal fibrosis in DDC-treated control livers, which was significantly attenuated in Alb-TEAD1^-/-^ mice (Figure 3A). Consistent with these findings, immunoblot analysis revealed strong induction of αSMA, a marker of activated stellate cells, in control livers following DDC injury, whereas αSMA expression was reduced in TEAD1-deficient livers (Figure 3B). To further assess stromal activation, we evaluated proliferating fibroblastic cells by co-staining for αSMA and Ki67. DDC injury led to increased αSMA^+^Ki67^+^ cells in control livers, indicative of active fibrogenic expansion, which was significantly reduced in Alb-TEAD1^-/-^ mice (Figure 3C). At the transcriptional level, quantitative PCR analysis demonstrated robust induction of profibrotic genes, including *Ctgf, Cyr61*, and *Spp1 (Osteopontin)*, in control livers following DDC exposure. Expression of these genes was consistently reduced in Alb-TEAD1^-/-^ livers (Figure 3D-F), supporting a role for hepatocyte TEAD1 in regulating fibrogenic signaling programs during injury. Together, these data indicate that hepatocyte TEAD1 contributes to both structural and transcriptional features of fibrogenic remodeling.

**Figure 3.**
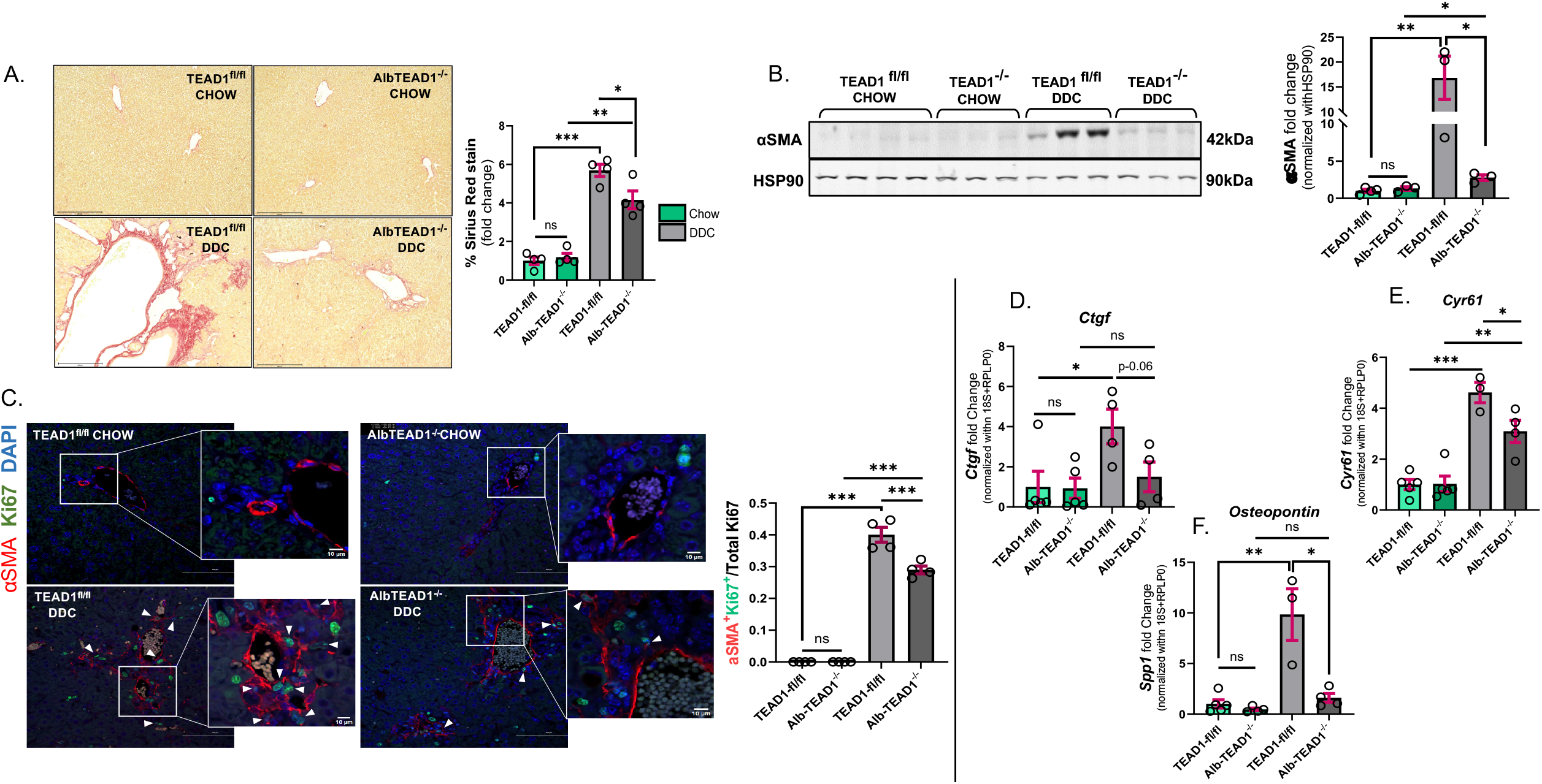
Decreased collagen deposition and profibrotic signaling in TEAD1-deficient livers. **(A)** Representative Picro-Sirius Red–stained liver sections and quantification of collagen-positive area from chow and DDC-fed TEAD1^fl/fl^ and Alb-TEAD1^-/-^ mice. **(B)** Representative immunoblot of αSMA from 30μg liver lysates. HSP90 was used as an endogenous control. **(C)** Representative immunofluorescence images and quantification of αSMA^+^Ki67^+^ cells with αSMA (red) and Ki67 (green) with DAPI (blue) signals. **(D-F)** Relative mRNA expression of profibrotic genes *Ctgf, Cyr61*, and *Spp1*. Data are presented as mean ± SEM (n = 3-5 mice per group). Images were taken at 20X and insets show magnified view, scale bar = 10 µm. [Statistical significance was determined by one-way ANOVA with post hoc testing (*p ≤ 0.05, **p ≤ 0.01, ***p ≤ 0.001)].

### Hepatocyte TEAD1 deficiency attenuates inflammatory cell recruitment during cholestatic injury

Inflammatory cell recruitment is a key driver of fibrosis progression in cholestatic liver disease. Immunostaining for F4/80 demonstrated marked macrophage accumulation in control livers following DDC treatment, whereas macrophage infiltration was reduced in Alb-TEAD1^-/-^ livers (Figure 4A). This was supported by decreased expression of macrophage-associated genes, including *Adgre1* and *Vcam1*, in TEAD1-deficient livers (Figure 4B-C). In addition to macrophages, neutrophil infiltration was assessed by Ly6G staining. DDC-treated control livers exhibited increased Ly6G^+^ cells, whereas Alb-TEAD1^-/-^ livers showed a reduction in neutrophil accumulation (Figure 4D). Consistent with reduced inflammatory recruitment, expression of the pro-inflammatory cytokine Tnfα was significantly lower in TEAD1-deficient livers compared with controls (Figure 4E). These findings suggest that hepatocyte TEAD1 contributes to the inflammatory milieu during cholestatic injury, potentially linking epithelial injury to immune cell recruitment.

**Figure 4.**
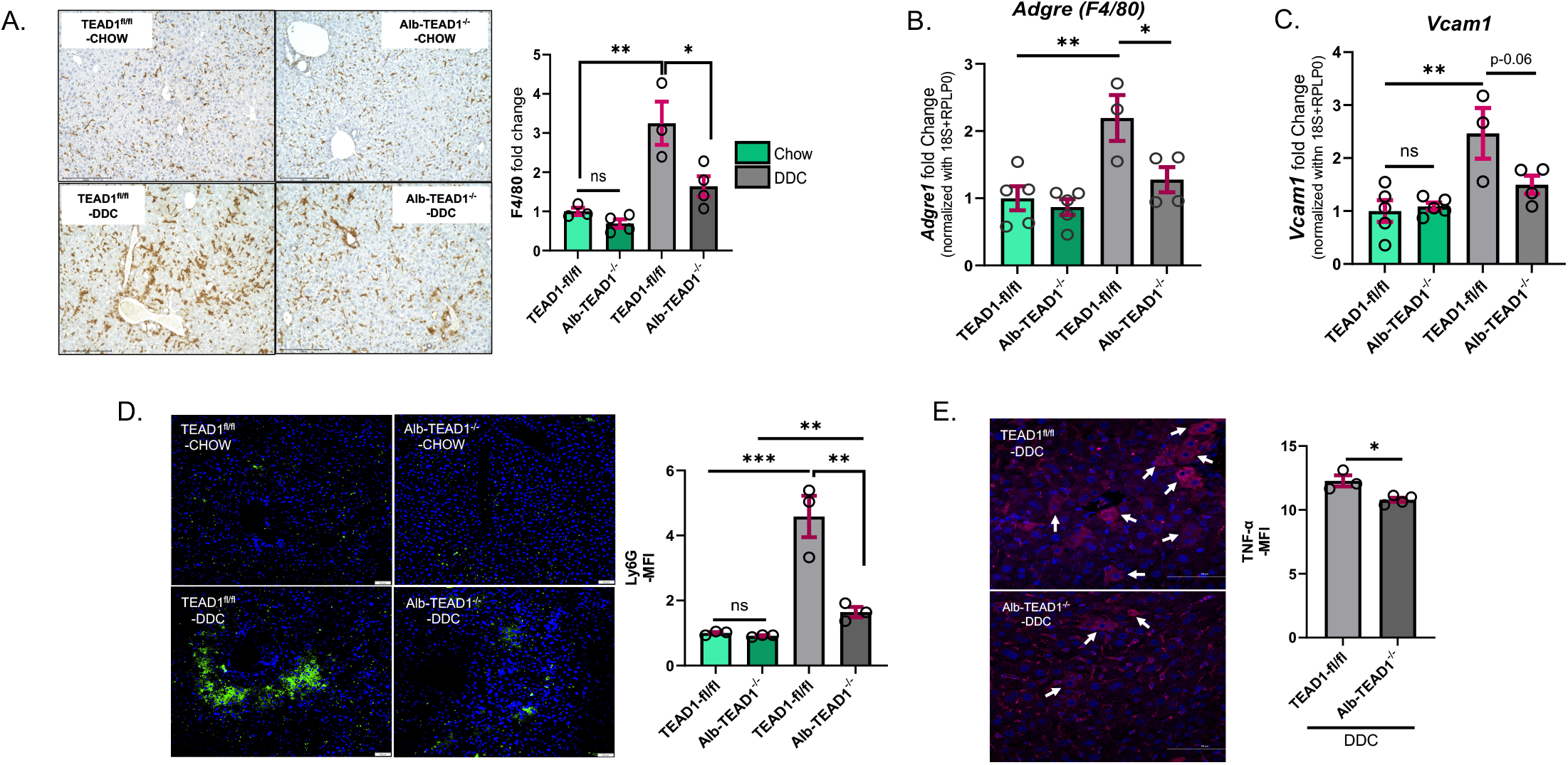
Reduced hepatic immune cell accumulation in Alb-TEAD1^-/-^ mice during cholestatic injury. **(A)** Representative immunohistochemistry images and quantification of F4/80^+^ macrophages in liver sections. **(B-C)** Relative mRNA expression of macrophage-associated genes *Adgre1* and *Vcam1*. Geomean of Ct values of 18S and RPLP0 was used as an endogenous control **(D)** Representative immunofluorescence images and quantification of Ly6G^+^ neutrophils. **(E)** Representative immunofluorescence images of Tnfα. Data are presented as mean ± SEM (n = 3-5 mice per group). Images were taken at 20X. [Statistical significance was determined by one-way ANOVA with post hoc testing (*p ≤ 0.05, **p ≤ 0.01, ***p ≤ 0.001)]. MFI-Mean Fluorescence Intensity.

### Hepatocyte TEAD1 deficiency reduces fibrosis without impairing hepatocyte proliferative responses

To determine whether reduced fibrosis in Alb-TEAD1^-/-^ livers was associated with impaired hepatocyte regeneration, we assessed proliferation using BrdU incorporation. DDC treatment induced hepatocyte proliferation in control livers, as expected; notably, Alb-TEAD1^-/-^ livers exhibited a modestly increased number of BrdU^+^ hepatocytes (Figure 5). These data indicate that hepatocyte-specific TEAD1 deletion does not compromise regenerative responses following injury. Instead, TEAD1 deficiency appears to selectively attenuate fibrogenic and inflammatory remodeling without impairing hepatocyte proliferation.

**Figure 5.**
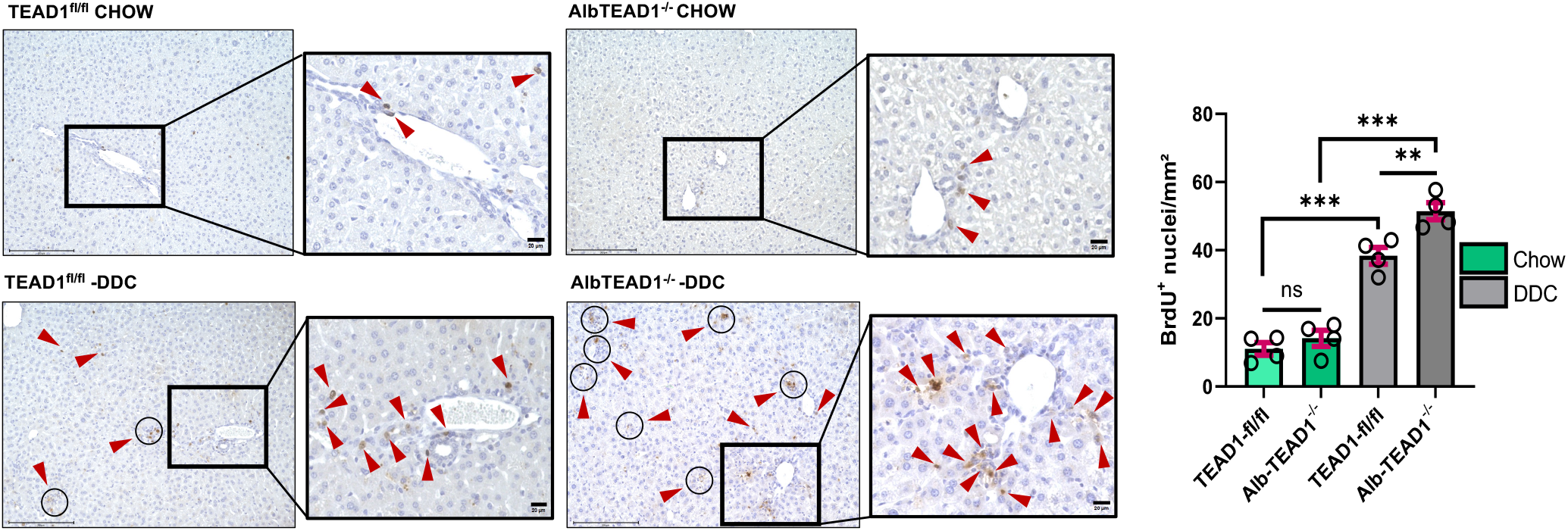
Preserved hepatocyte proliferation in TEAD1-deficient livers following DDC injury. Representative images of BrdU staining and quantification of BrdU^+^ cells in liver sections from DDC-treated mice. Data are presented as mean ± SEM (n = 3-5 mice per group). Images were taken at 20X and insets show magnified view, scale bar = 10 µm. [Statistical significance was determined by one-way ANOVA with post hoc testing (**p ≤ 0.01, ***p ≤ 0.001)].

### Bile acid transport and synthetic responses to cholestatic injury are preserved in the absence of hepatocyte TEAD1

To determine whether hepatocyte TEAD1 influences bile acid homeostasis during cholestatic injury, we examined the expression of genes involved in hepatic bile acid transport and synthesis. As expected, DDC treatment suppressed the uptake transporters *Oatp1, Oatp4*, and *Ntcp*, consistent with adaptive responses to limit intracellular bile acid accumulation. These changes were comparable between TEAD1^fl/fl^ and Alb-TEAD1^-/-^ livers (Figure 6A-C). DDC exposure also induced the efflux transporters *Mrp2* and *Mrp3*, along with increased expression of *Abcb4* and *Slc51b*. While modest differences were observed in Alb-TEAD1^-/-^ livers, these did not reach statistical significance, and overall induction patterns were similar between genotypes (Figure 6D-G). In parallel, DDC suppressed the bile acid synthetic enzymes *Cyp7a1, Hsd3b7, Cyp8b1*, and *Cyp7b1* of both classical and alternative pathways of bile acid synthesis. Although Alb-TEAD1^-/-^ mice exhibited slightly elevated basal expression of select genes under chow conditions, repression of the bile acid synthetic program following DDC exposure was comparable between groups (Figure 6H-K). Collectively, these findings indicate that hepatocyte TEAD1 does not substantially alter canonical transcriptional responses governing bile acid transport and synthesis during cholestatic injury.

**Figure 6.**
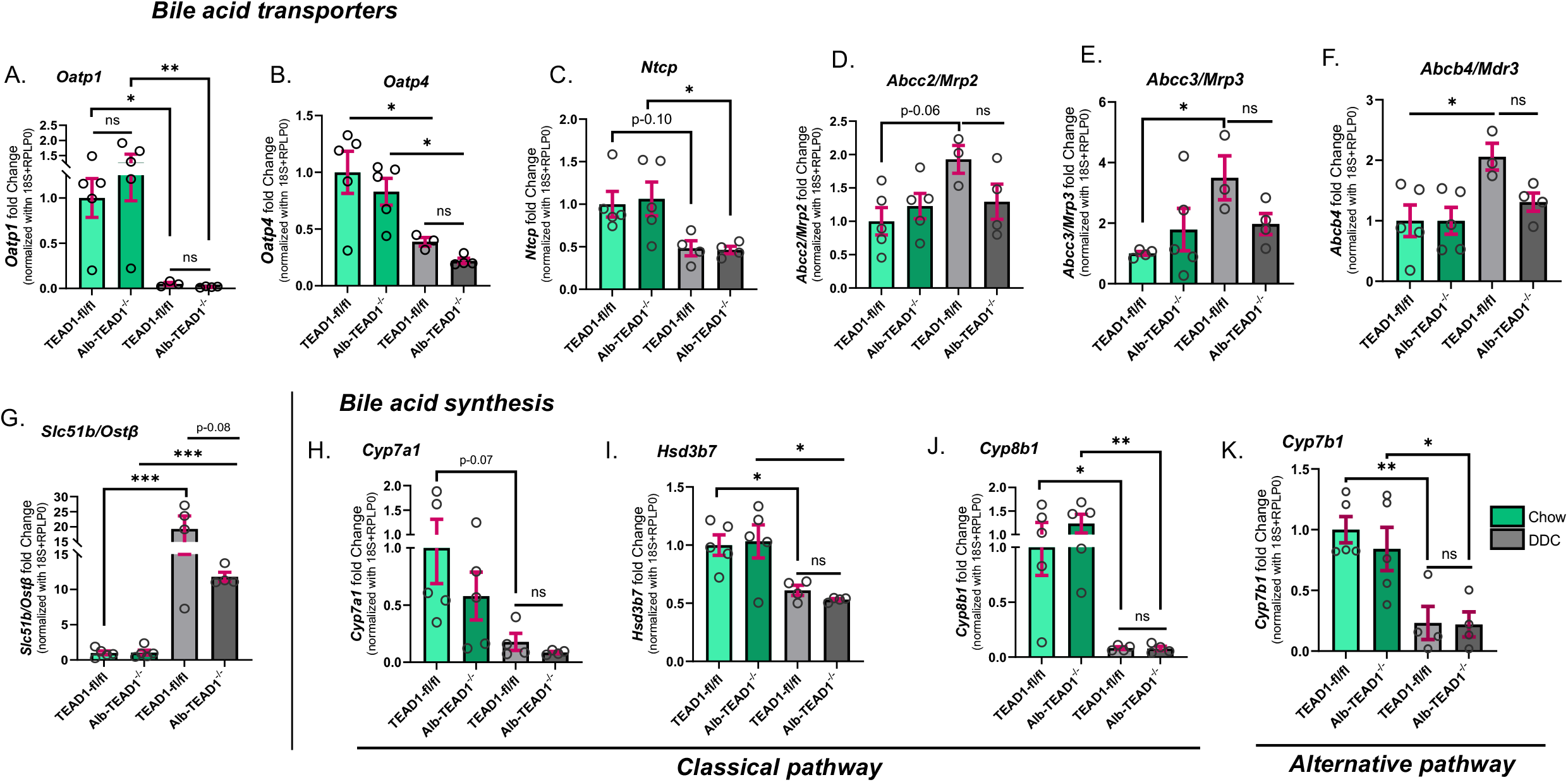
Comparable regulation of bile acid transport and synthesis genes in control and TEAD1-deficient livers. **(A-C)** Relative mRNA expression of bile acid uptake transporters *Oatp1, Oatp4*, and *Ntcp*. **(D-G)** Relative mRNA expression of bile acid efflux transporters *Mrp2, Mrp3, Abcb4*, and *Slc51b*. **(H-K)** Relative mRNA expression of bile acid synthetic enzymes *Cyp7a1, Hsd3b7, Cyp8b1*, and *Cyp7b1*. Geomean of Ct values of 18S and RPLP0 was used as an endogenous control. Data are presented as mean ± SEM (n = 3-5 mice per group). [Statistical significance was determined by one-way ANOVA with post hoc testing (*p ≤ 0.05, **p ≤ 0.01, ***p ≤ 0.001)].

### TEAD1-associated fibrogenic programs are elevated in human PSC liver samples

To evaluate the translational relevance of our findings, we analyzed publicly available transcriptomic data from human PSC liver samples (GSE159676). Expression of TEAD1 and TEAD1-associated profibrotic genes, including *CTGF, SPP1, ANKRD1*, and *AXL* were significantly increased in PSC livers compared with healthy controls (Figure 7A-E). While these analyses do not establish causality, they are consistent with our experimental findings and support a potential role for TEAD1-associated transcriptional programs in human cholestatic liver disease.

**Figure 7.**
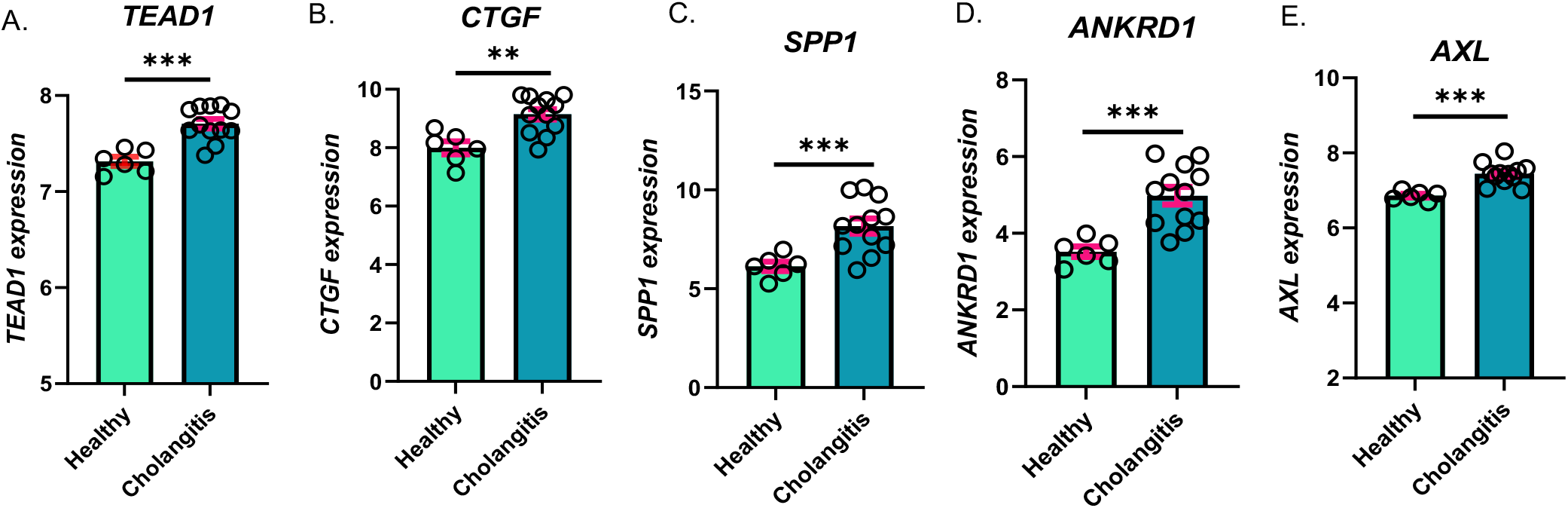
Elevated TEAD1-associated gene expression in human PSC liver samples. **(A-E)** Relative gene expression (normalized log_2_ values) of TEAD1 and TEAD-associated targets (*CTGF, SPP1, ANKRD1, AXL*) in liver samples from healthy controls and patients with primary sclerosing cholangitis (PSC) (GSE159676). Data are presented as mean ± SEM. [Statistical significance was determined using an unpaired Welch’s t-test (**p ≤ 0.01, ***p ≤ 0.001)].

### Hepatocyte-enriched TEAD1 and profibrotic co-expression are upregulated in single-cell liver transcriptome of human PSC patients

To extend our translational analysis at single-cell resolution, we interrogated a published human PSC liver single-cell atlas comprising hepatocytes, immune cells, endothelial populations, and stromal cells from PSC patients and non-diseased donors (NDD) [7]. Dot-plot analysis across all cell types demonstrated that TEAD1 is broadly expressed in the PSC liver, with the highest expression detected in all three hepatocyte zones (periportal, midzonal, and centrilobular region hepatocytes) (Figure 8A). Among TEAD1 target genes, CCN1 (CYR61) and CCN2 (CTGF) expression was most prominent in hepatic stellate cells, whereas SPP1 and ANKRD1 showed enrichment in cholangiocytes, and AXL was highly expressed in fibroblasts, Kupffer cells, and macrophages, with monocytes and stellate cells displaying moderate expression (Figure 8A). UMAP embedding and violin plot analysis confirmed that TEAD1 expression was increased in PSC compared with NDD livers across multiple cell types, with the most pronounced differences observed in hepatocyte clusters (Figure 8B).

**Figure 8.**
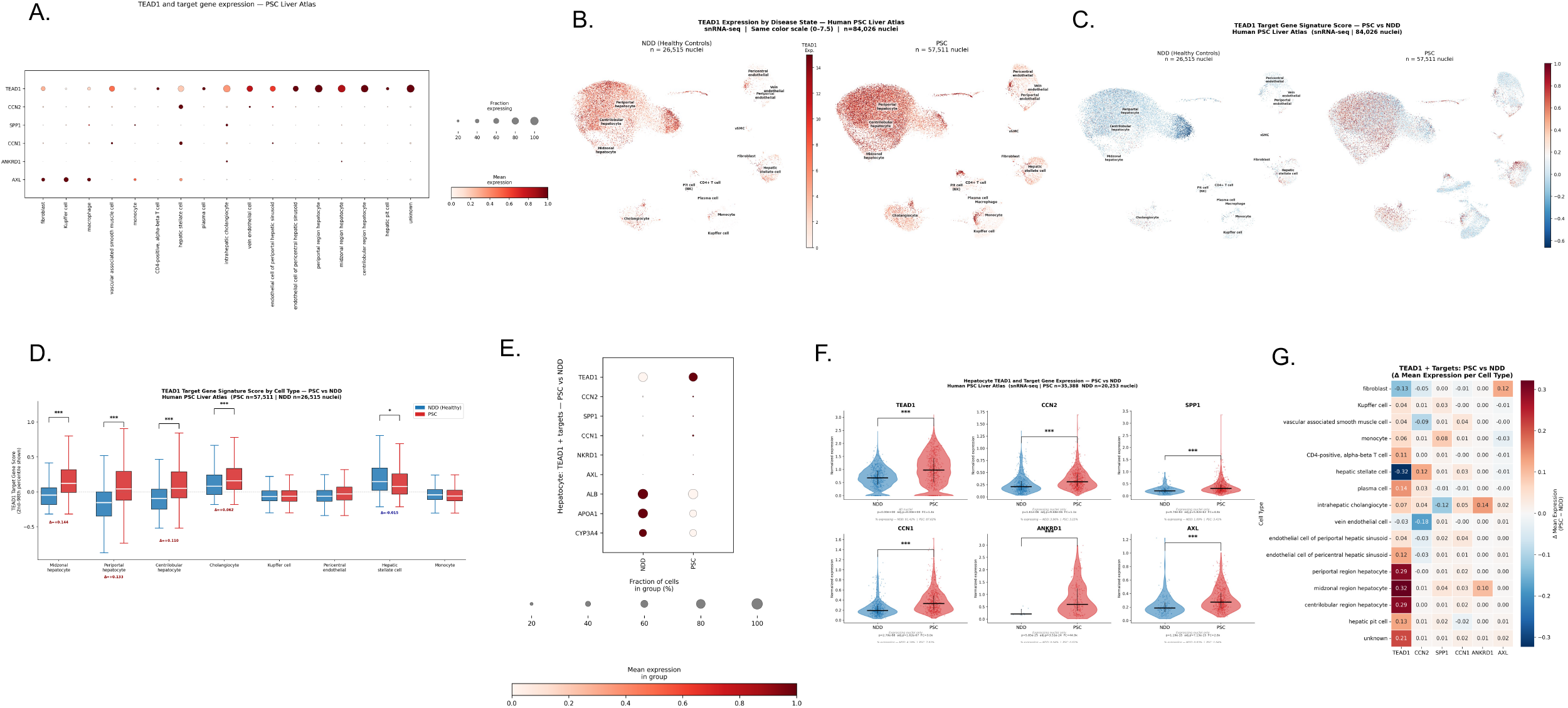
TEAD1 and target gene expression in human PSC liver at single-nucleus resolution. snRNA-seq analysis of human PSC liver atlas (GSE245620; PSC n = 7 donors, 57,511 nuclei; healthy controls [NDD] n = 3 donors, 26,515 nuclei; total n = 84,026 nuclei). **(A)** Dot plot of TEAD1 and target gene expression across all annotated liver cell types (PSC + NDD combined). Dot size = fraction of nuclei expressing; dot color = mean normalized expression (scaled per gene). **(B)** UMAP of TEAD1 expression in NDD (left) and PSC (right) nuclei. The color scale (0–7.5) indicates expression. Cell type annotations shown at cluster centroids. **(C)** UMAP of TEAD1 target gene module score in NDD (left) and PSC (right). Score computed by Scanpy score_genes; positive values (red) indicate above-expected TEAD1 target activity. Identical diverging color scale for both panels. **(D)** Box plots of TEAD1 target gene module scores by cell type, PSC (red) vs NDD (blue), ordered by effect size (Δ = PSC − NDD mean). Two-sided Mann-Whitney U test, Bonferroni correction. ***p < 0.001; *p < 0.05. **(E)** Dot plot of TEAD1 target genes and hepatocyte identity markers in hepatocytes, PSC vs NDD. **(F)** Violin plots of individual gene expression in hepatocytes, PSC vs NDD. % expressing indicated below each panel. Horizontal bar = median; vertical line = IQR. All adj.p < 0.001 (Bonferroni, n = 6 comparisons). **(G)** Heatmap of Δ mean expression (PSC − NDD) for TEAD1 and target genes across all cell types.

To assess TEAD1 target activation at the individual cellular level, we computed a composite TEAD1 target gene signature score across cell types (see Methods). UMAP visualization revealed that the highest signature scores clustered predominantly within hepatocyte populations and within cholangiocyte, Kupffer, and stellate cell clusters (Figure 8C). Quantitative comparison of signature scores between PSC and control NDD demonstrated that the TEAD1 target gene program was significantly elevated in all three hepatocyte zones (periportal: delta = 0.133, p-adj approximately 0; centrilobular: delta = 0.110, p-adj approximately 0; midzonal: delta = 0.144, p-adj = 3.25.x10^-31^) in PSC relative to NDD, with effect sizes substantially larger than those observed in non-parenchymal populations (Figure 8D). Consistent with the mouse data, dotplot analysis of PSC versus NDD hepatocytes demonstrated concurrent TEAD1 and its targets’ upregulation alongside downregulation of canonical hepatocyte function markers (ALB, APOA1, and CYP3A4), suggesting that TEAD1 activation in PSC hepatocytes is accompanied by a partial loss of differentiated hepatocyte identity (Figure 8E). Violin plot analysis confirmed that expression of individual TEAD1 target genes was significantly increased in PSC hepatocytes relative to controls (all adj. p < 0.001), supporting activation of a TEAD1-dependent transcriptional program at the single-cell level (Figure 8F). Differential expression heatmap analysis further showed that TEAD1 upregulation was most pronounced in hepatocyte populations, with midzonal (delta = 0.32) and periportal/centrilobular region hepatocytes (delta = 0.29 each) displaying the largest increases across all cell types examined (Figure 8G).

Collectively, these human PSC single-cell analyses corroborate the mouse DDC-diet findings and identify hepatocyte TEAD1 as a reproducibly elevated, transcriptionally active signal in human PSC liver.

## DISCUSSION

Cholestatic liver injury is characterized by impaired hepatocyte regeneration, persistent ductular signaling, and progressive fibrogenic remodeling. In this study, we identify hepatocyte TEAD1 as a context-dependent transcriptional regulator that amplifies injury-associated ductular, inflammatory, and fibrogenic responses without substantially altering core bile acid metabolic adaptation. Hepatocyte-specific deletion of TEAD1 attenuated DDC-induced fibrosis, inflammatory cell recruitment, and ductular expansion, while preserving hepatocyte proliferative capacity during active injury. These findings support a model in which hepatocyte TEAD1 contributes to the selection of a fibrogenic repair trajectory during cholestatic stress.

The Hippo-YAP-TEAD axis has been extensively implicated in liver regeneration, fibrosis, and tumorigenesis. Prior studies have primarily focused on TEAD1 function in non-parenchymal compartments, particularly hepatic stellate cells, where TEAD1 cooperates with YAP/TAZ to promote extracellular matrix production and cellular activation [5, 6]. Consistent with this framework, we observed robust induction of canonical TEAD target genes, including *Ctgf, Cyr61*, and *Spp1*, in DDC-treated livers. Importantly, liver-specific deletion of TEAD1 significantly reduced the expression of these genes, suggesting that hepatocyte-intrinsic TEAD1 activity contributes to the establishment of a profibrotic transcriptional program and shaping the fibrogenic microenvironment during cholestatic injury.

Ductular reaction is a defining feature of PSC and other cholestatic disorders and is closely associated with fibrosis progression and clinical outcome. In this study, TEAD1 deficiency led to a marked reduction in CK19^+^ ductular structures and intraductal remodeling following DDC exposure. Although the precise cellular origin and plasticity of ductular cells remain under investigation, our findings suggest that hepatocyte-derived transcriptional programs contribute to ductular remodeling. In this context, TEAD1 may function upstream of epithelial-stromal signaling networks that coordinate biliary expansion and fibrogenesis.

Inflammatory attenuation in TEAD1-deficient livers paralleled suppression of ductular and fibrogenic responses. Reduced macrophage and neutrophil accumulation, together with decreased expression of inflammatory mediators such as *Vcam1* and TNFα, indicates that hepatocyte TEAD1 contributes to the establishment of a pro-inflammatory microenvironment during cholestatic injury. Given the known interplay between inflammatory signaling and fibrogenesis, it is likely that these effects are interdependent. The concurrent suppression of *Spp1*, a mediator implicated in both immune cell recruitment and ductular expansion, further supports a model in which hepatocyte TEAD1 amplifies epithelial-derived inflammatory signals that reinforce fibrogenic progression.

A notable finding of this study is that hepatocyte-specific TEAD1 deletion attenuated fibrosis and inflammation without impairing hepatocyte proliferative responses. BrdU incorporation demonstrated that hepatocyte proliferation was preserved, and in some cases modestly enhanced, in TEAD1-deficient livers following injury. This suggests that TEAD1 deficiency selectively limits maladaptive remodeling while maintaining regenerative capacity. This observation is consistent with prior work demonstrating that TEAD1 can restrain proliferation in differentiated cell types, including pancreatic β cells [8]. However, whether similar transcriptional mechanisms operate in hepatocytes remains to be determined.

Importantly, hepatocyte TEAD1 deficiency did not substantially alter transcriptional programs governing bile acid transport or synthesis. Canonical adaptive responses to DDC-induced cholestasis, including suppression of uptake transporters and bile acid synthesis genes and induction of efflux pathways, were preserved in TEAD1-deficient livers. These findings suggest that the protective effects of TEAD1 deletion are not mediated by restoring bile acid homeostasis but instead reflect selective modulation of injury-associated fibrogenic and inflammatory pathways. This distinction positions TEAD1 downstream or independent of primary cholestatic sensing mechanisms.

The translational relevance of these findings is supported by analysis of human PSC transcriptomic datasets [9], which demonstrate coordinated upregulation of TEAD1 and TEAD1-associated targets, including *CYR61, ANKRD1, CTGF, AXL*, and *SPP1*. Importantly, single-cell transcriptomic analyses further refine this observation, revealing hepatocyte-enriched TEAD1 expression and activation of a TEAD1-dependent transcriptional program across all hepatic zones in PSC. While these data do not establish causality, the convergence of bulk and single-cell evidence with our hepatocyte-specific genetic model supports TEAD1 activation as a conserved feature of cholangiopathy-associated fibrosis. Prior studies have linked Hippo pathway dysregulation to advanced liver disease [10]; our findings extend these observations by providing direct hepatocyte-specific genetic evidence implicating TEAD1 as a regulator of fibrogenic remodeling in cholestatic injury. Together, these data suggest that TEAD1 activation reflects a maladaptive hepatocyte state that contributes to epithelial-stromal crosstalk, rather than a secondary consequence of injury, and highlight TEAD1-directed signaling as a potential therapeutic axis in PSC.

Several limitations should be considered. The DDC model recapitulates key features of ductular and periportal fibrogenic remodeling but does not fully capture the chronic and heterogeneous progression of human PSC. Long-term and complementary injury models will be required to determine whether sustained TEAD1 suppression confers durable antifibrotic benefit. In addition, while our genetic model defines a hepatocyte-intrinsic role for TEAD1, the contributions of TEAD signaling in non-parenchymal cells, including cholangiocytes, immune populations, and fibroblasts, remain to be delineated. Finally, the direct transcriptional targets of TEAD1 in hepatocytes during cholestatic injury have not been defined and will require future chromatin-based analyses.

From a therapeutic perspective, the YAP-TEAD axis represents a pharmacologically attractive target. Emerging TEAD inhibitors, including palmitoylation inhibitors such as VT3989, as well as agents that disrupt YAP-TEAD interactions, provide opportunities to test whether modulation of this pathway can selectively attenuate fibrogenic remodeling while preserving regenerative responses. In this context, human-relevant systems, including liver-on-chip platforms and hepatocyte-stromal co-culture models, may be particularly informative for defining the therapeutic window and cell-type specificity of TEAD-directed interventions prior to translation into clinical trials.

In summary, our findings identify hepatocyte TEAD1 as a regulator of epithelial-stromal remodeling during cholestatic liver injury. By selectively promoting ductular expansion, inflammatory recruitment, and fibrogenic signaling while sparing core metabolic adaptation and regenerative capacity, hepatocyte TEAD1 emerges as a potential target for uncoupling maladaptive fibrosis from essential tissue repair processes in cholestatic liver disease.

## MATERIAL AND METHODS

### Animals used in the study

Hepatocyte-specific TEAD1 knockout (Alb-TEAD1^-/-^) mice were generated by crossing TEAD1^fl/fl^ mice with transgenic mice expressing Cre recombinase under the control of the albumin (Alb) promoter. Male C57BL/6 mice aged 6-8 weeks were used for all experiments. Mice were randomly assigned to four groups: TEAD1^fl/fl^ chow (n = 5), Alb-TEAD1^-/-^ chow (n = 5), TEAD1^fl/fl^ DDC (n = 4), and Alb-TEAD1^-/-^ DDC (n = 4). For induction of cholestatic injury, mice were fed a standard chow diet supplemented with 0.1% (w/w) 3,5-diethoxycarbonyl-1,4-dihydrocollidine (DDC; Sigma-Aldrich) for 8 consecutive days. For proliferation assays, 5-bromo-2′-deoxyuridine (BrdU; Sigma-Aldrich, B9285) was administered intraperitoneally (15 μL/g body weight) 2 hours prior to sacrifice. All animals were maintained under specific pathogen-free conditions in the University of Pittsburgh Division of Laboratory Animal Resources facility, with controlled temperature and humidity, a 12-hour light/dark cycle, and ad libitum access to food and water.

### Sample Isolation

Livers from these mice were isolated and immediately shock frozen in liquid nitrogen and stored at -80 degrees, until next use for RNA and protein isolation. Livers were also soaked in paraformaldehyde for paraffin embedding and sectioning.

### Histology and tissue fibrosis

Paraffin embedded tissues were sectioned to 5 µm thickness. Tissue histology was accessed by hematoxylin and eosin (H&E) kit (Abcam, ab245880) and liver fibrosis was accessed with Picro Sirius Red staining kit (Abcam, ab150681), as per the manufacturer’s protocol.

### Immunostaining

Paraffin-embedded liver sections were deparaffinized, rehydrated through graded ethanol, and subjected to antigen retrieval, followed by permeabilization. Sections were incubated with primary antibodies overnight at 4°C and subsequently with appropriate secondary antibodies for 1 hour at room temperature as previously described [11]. For immunohistochemistry, signal detection was performed using the VECTASTAIN® Elite® ABC-HRP Kit (PK-6100, Vector Laboratories). For immunofluorescence, fluorophore-conjugated secondary antibodies (Alexa Fluor 488, 1:500; Alexa Fluor 546, 1:1000; Alexa Fluor 647, 1:500) were used.

Primary antibodies included F4/80 (1:250; Cell Signaling Technology, 70076S), CK19 (1:200; Novus, NB100-687SS), BrdU (1:200; Abcam, ab6326), Albumin (1:500; Abcam, ab19194), TNFα (1:250; Abcam, ab1793), TEAD1 (1:200; Abcam, ab133533), Ly6G (1:250; Santa Cruz, sc-53515), αSMA (1:500; Invitrogen, 14-9760-82), and Ki67 (1:500; Cell Signaling Technology, 9129). Images were acquired using an Olympus AX70 microscope and an all-in-one APX100 system and processed with cellSens software; confocal imaging was performed on a Nikon A1 system. IHC images were acquired on EVOS FL Auto 2 system. Signal quantification was performed using ImageJ software.

### Gene and protein expression

#### qPCR

Total RNA was isolated from liver tissue and treated with DNase to remove genomic DNA contamination. cDNA was synthesized from 500 ng of RNA using standard reverse transcription methods. Quantitative real-time PCR (qRT-PCR) was performed using gene-specific primers and 2X SYBR Green Master Mix (GenDEPOT, Q5600-005) on a QuantStudio 5 Real-Time PCR System (Applied Biosystems). Gene expression levels were normalized to housekeeping genes (geometric mean of 18S and RPLP0) and calculated using the comparative Ct (ΔΔCt) method. Primer sequences are provided in Table 1.

**TABLE 1.**
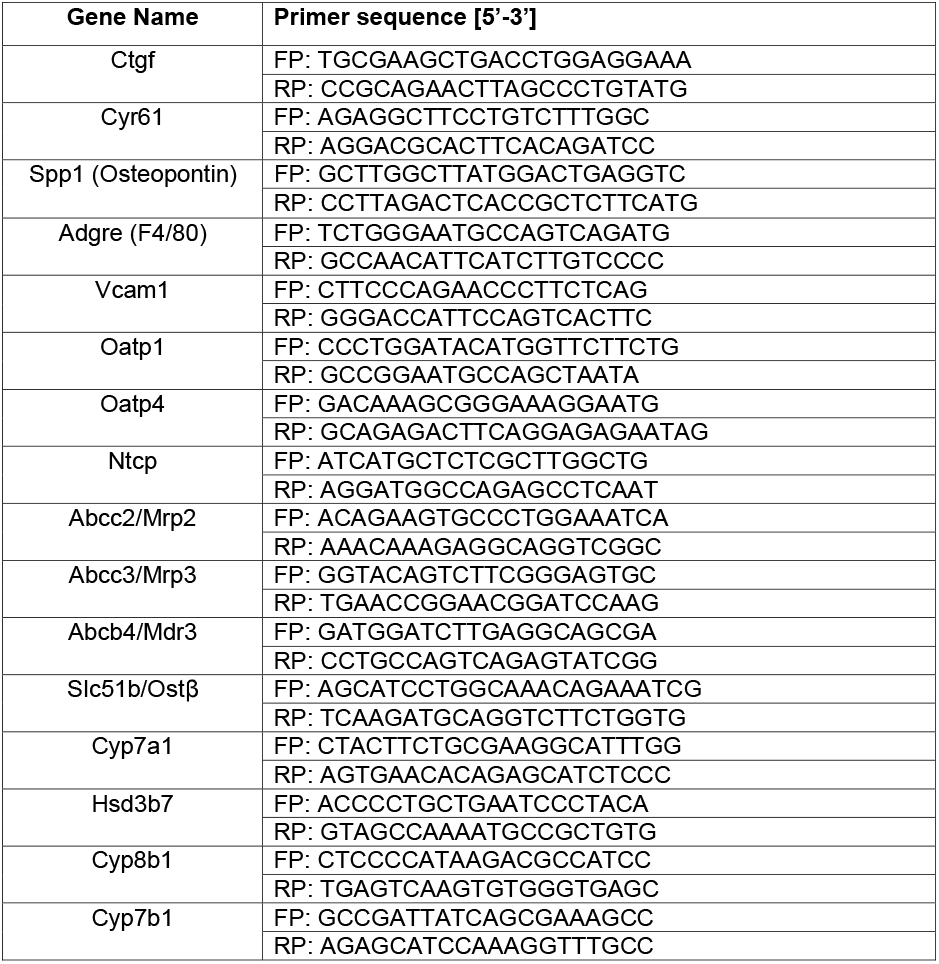
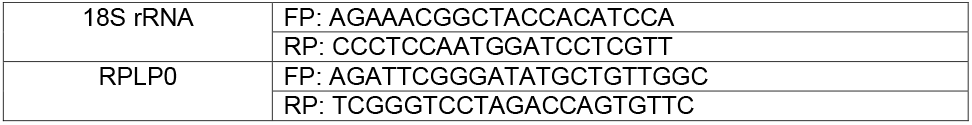
Primer sequence of genes analyzed.

#### Liver transcriptomic data

Publicly available liver transcriptomic data were obtained from the Gene Expression Omnibus (GEO) dataset GSE159676, comprising liver samples from healthy controls (n = 6) and patients with primary sclerosing cholangitis (PSC) (n = 12) [9]. Processed expression matrices were downloaded directly from GEO. Expression values were generated using Affymetrix Human Gene 1.0 ST arrays and had undergone background correction, quantile normalization, and log_2_ transformation using the Robust Multi-array Average (RMA) method prior to deposition. Gene expression values for selected targets were extracted and compared between control and PSC samples.

#### Western blot

Snap frozen livers were digested and sonicated in the tissue lysis buffer supplemented with protease and phosphatase inhibitors and 30µg protein was separated through SDS-PAGE and transferred to nitrocellulose membrane [12, 13]. Primary antibody incubation was performed overnight at 4°C using the following antibodies: TEAD1 (Abcam-ab133533; 1:2000), αSMA (Invitrogen-14-9760-82; 1:1000). HSP90 (Santa Cruz-sc13119; 1:5000) was used as endogenous control. Secondary antibody incubation (1:5000) was performed for 2h at room temperature before signal acquisition.

#### Single-Nucleus RNA-seq Analysis

We analyzed a publicly available snRNA-seq dataset of human PSC and healthy control liver (GEO: GSE245620 [7]; PSC: n = 7 donors, 57,511 nuclei; healthy controls (no disease donors -NDD): n = 3 donors, 26,515 nuclei; total n = 84,026 nuclei; 10x Genomics Chromium 3’ v3), accessed via CellxGene (ID: 4b5895d7-6d92-471a-b13a-5c59a000ddc4). Pre-computed log-normalized expression values, cell type annotations, and UMAP embeddings were used without modification. A composite TEAD1 target gene module score was computed for each nucleus using Scanpy’s score genes function, representing the mean expression of 16 TEAD1 target genes (TEAD1, CCN2, SPP1, CCN1, ANKRD1, AXL, AMOTL2, AREG, BIRC5, CCND1, ITGB2, LATS2, MYC, PTPN14, TGFB1, THBS1, VGLL4) minus the mean of expression-matched control genes (50 controls per gene; 25 bins). Differential expression between PSC and NDD nuclei was assessed by two-sided Mann-Whitney U test with Bonferroni correction. Hepatocyte analyses combined periportal, centrilobular, and midzonal hepatocyte nuclei (PSC n = 35,388; NDD n = 20,253). For sparsely expressed genes (< 8% expressing), violin plots show only nuclei with detectable expression (> 0); the horizontal bar and vertical line within each violin represent the median and interquartile range respectively. Pearson correlations between TEAD1 and target genes were computed separately in PSC and NDD nuclei expressing both genes simultaneously. Analyses used Python 3.10, Scanpy 1.11.5, and SciPy 1.15.3.

### Statistical analysis

Statistical analyses were performed using GraphPad Prism (version 9.0). Comparisons between two groups were conducted using an unpaired Welch’s t-test, while multiple group comparisons were analyzed by one-way ANOVA with appropriate post hoc testing. A p value ≤ 0.05 was considered statistically significant.

## Authorship Contributions

Conceptualization and study design (VY, AK, JKL). Data generation (AK, JKL, VY). Single cell data analysis (VY, AK). Data analysis and interpretation (AK, VY, JKL, VN, VM, DF, JD, RP, SG, MM). Image acquisition (RP, SG, AK). Manuscript preparation (AK,VY). Critical review and editing (AK, VY, MM). Funding-(VY).

## Data Availability

The Data in this study will be made available upon reasonable request from the corresponding author.

